# An RF coil array system for in vivo & ex vivo non-human primate brain studies at 10.5 Tesla

**DOI:** 10.64898/2025.12.09.692206

**Authors:** Matt Waks, Alexander Bratch, Steve Jungst, Russell L. Lagore, Benjamin C. Tendler, Shaun Warrington, Wenchuan Wu, Stamatios N. Sotiropoulos, Karla L. Miller, Steen Moeller, Ana M.G. Manea, Kamil Uğurbil, Jan Zimmermann, Gregor Adriany

**Author notes:** Authors contributed equally to this work (Matt Waks, Alexander Bratch). Authors contributed equally to this work (Jan Zimmermann, Gregor Adriany). **CORRESPONDING AUTHOR:** Matt Waks Center for Magnetic Resonance Research (CMRR) University of Minnesota 2021 6th Street SE, Minneapolis, MN 55455.

## Abstract

**Purpose:** We aimed to develop an innovative RF coil toolset for high resolution in vivo and ex vivo non-human primate neuroimaging applications at 10.5T MRI. The main goal was to improve our multimodal neural connectivity pipeline through the improved SNR, CNR, and spatial resolution that result from utilizing high-density RF coil arrays in combination with ultra-high field MRI.

**Methods:** Two RF coil arrays, both comprising 8 transmit and 40 receive channels, were designed and built for in vivo and ex vivo NHP imaging at 10.5 T. Experimental data was collected on multiple in vivo and ex vivo specimens, showing the utility UHF MRI for NHP neuroscience.

**Results:** Experimental results demonstrate uniform whole-brain coverage within both in vivo—which included the cerebellar, and spinal cord regions—and ex vivo specimens. High spatial resolutions *(in vivo: 0.1×0.1×0.5 mm, ex vivo: 120 micron isotropic*) were achieved, revealing detailed anatomical structures throughout. High temporal stability allowed for EPI and dMRI applications.

**Conclusion:** An RF coil toolset comprising two 8ch transmit/40ch receive arrays was successfully designed for both in vivo and ex vivo NHP brain applications at 10.5 T. Both high-density RF coil arrays coupled with the SNR and CNR benefits available at UHF supported unmatched anatomical, functional, and diffusion applications at some of the highest spatial resolutions to date. Additionally, the combined capability of both in vivo and ex vivo MRI with the same brain specimen allows for tighter control for experimental-paradigm multistage studies examining translation between in vivo and ex vivo methodologies.

## 1 INTRODUCTION

Ultra-high field (UHF) MRI has given rise to whole-brain mesoscopic-scale imaging with unmatched spatial and temporal resolution due to the improved signal-to-noise (SNR) and contrast-to-noise (CNR) that comes with higher field strength.^1^ As a result, UHF imaging is essential for the optimal visualization of fine details within deep brain structures, the distinction between cortical laminae, and the definition of axonal-level structures such as the line of Gennari. In addition to the intrinsic benefits in SNR and CNR afforded by UHF MRI, optimization of Radio Frequency (RF) coils has a considerable impact on the achievable SNR. Consequently, considerable effort has gone into the development of novel hardware in recent years, particularly in the form of high channel-count RF coil arrays. Coupled with parallel transmit technology to counter UHF wavelength effects, these arrays have been essential for advanced imaging techniques such as DWI, DTI, BOLD, and others to flourish, producing stunning results^2,3^ and revealing the functional pathways of the human brain.

At UHF—and for neuroscience applications in general—non-human primates (NHP) in particular have great translational promise.^4–8^ Of those, the cynomolgus (Macaca fascicularis) and rhesus monkey (Macaca mulatta) are the mostly widely used NHP species for applied biomedical research due to the similarities in physiology, development, genetics, and social interaction.^9^ There has been a large body of work focusing on RF coil design for NHP brain research at various field strengths^10–19^ and under various experimental and technical conditions.^20–24^ In our previously published work by Lagore et al,^25^ we demonstrated submillimeter-resolution imaging and tractography capabilities with a parallel transmit, parallel receive coil array at 10.5T.^8,26^

In line with our goal to support both software and hardware advancements in NHP imaging at UHF, we present herein two RF coil ensembles that form a toolset for both in vivo & ex vivo experiments to further expand NHP neuroimaging development at 10.5T. The underlying array technology in both cases is informed by our prior 10.5T in vivo human head arrays^3^ but to address the specific demands of NHP applications we needed to develop novel technological solutions resulting in optimized 40 channel arrays for ex-vivo and in-vivo imaging.

## 2 METHODS

### 2.1 RF Coils

Two 8-channel transmit, 40-channel receive (8Tx/40Rx) coil assemblies were constructed, one for in vivo and another for ex vivo (Figure 1) studies; each consisted of two subassemblies: an 8-channel loop coil array with each element capable of transmit and receive functions (8Tx/Rx, i.e., 8-channel “transceiver” array) and a 32-channel receive-only array (32Rx). All coil housing components were designed and 3D printed using a polycarbonate filament (PolyMax™ PC, Polymaker, Changshu, China) on either an X1 Carbon (Bambu Lab, Shenzhen, China) or M2+ (AON3D, Montreal, Canada) printer; no surface finishes were applied to the coil housings prior to the coil assembly process.

**Figure 1:**
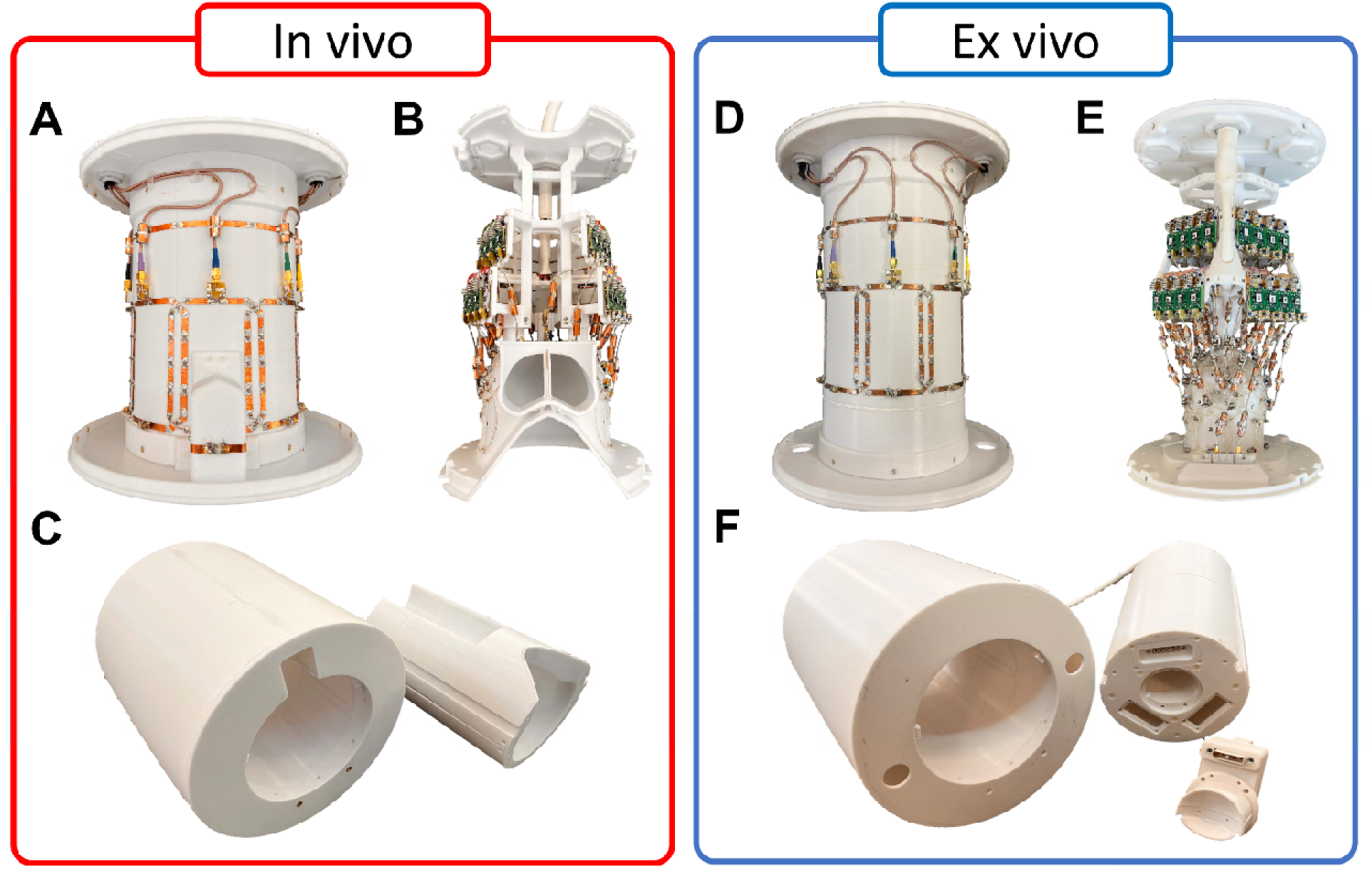
Subassemblies of the 8Tx/40Rx_i_ array: (A) 8Tx/Rx_i_ array, (B) 32Rx_i_ array, and (C) the complete coil ensemble. Subassemblies of the 8Tx/40Rx_e_ array: (D) 8Tx/Rx_e_ array, (E) 32Rx_e_ array, and (F) the complete coil ensemble.

The first of two coils supports in vivo applications of various macaque species. In order to harmonize both acquisition parameters and imaging artifacts for improved translation between NHP and human research, this coil was designed for head first supine orientation of the animal. Additionally, this coil was developed with fMRI applications in mind, and thus incorporates large openings at the anterior of the coil former used for visual stimulation and animal support. Due to the change in orientation, the larger anterior openings, and the intrinsic differences between an NHP and human head (shallower height, larger snout and temporalis muscles), consideration needed to be given to an array architecture that would not result in a loss of sensitivity in the areas associated with these changes (e.g., anterior and frontal regions). A novel implementation of a detunable, reactively-shortened dipole antenna was chosen to increase signal reception in these hard to capture regions of the brain.^27^

The second coil, supports ex vivo MRI applications within the same genus. As is typical of ex vivo neuro MRI, the brain is fully circumscribed by an array of receive coils. A novel feature of this design, is that the coil array is not split along the coronal plane, but rather was designed as a typical in vivo array with an opening at the inferior end, where we have now included a plug—comprising two receive-only coil elements—to be placed in the inferior opening, fully encapsulating the ex vivo brain.

Similar to our human studies at 10.5T,^3,28^ the use of the transmit array as additional receivers increases the SNR in both the in vivo and ex vivo arrays. This toolset was designed for imaging the same brain specimen throughout its active life and post-mortem. With the capability to collect data in this fashion, pre- and post-mortem, we present a toolset to further expand neuroscience applications. However, the results shown are from multiple specimens both in vivo and ex vivo.

#### 2.1.1 8-channel Transmit/Receive Arrays

Two 8Tx/Rx arrays (Figure 1A and D) were constructed (8Tx/Rx_i_ for in vivo, 8Tx/Rx_e_ for ex vivo), each consisting of one row of eight 9 x 12.5 cm coil elements mounted directly to a 19.6 cm diameter cylindrical former, and overlapped in the azimuthal direction to minimize magnetic coupling between one another. Each element was fabricated with ∼75 µm thick copper conductors and trace widths of 6.5 mm. The transceiver housing, designed to be compatible with a head gradient insert coil, was 41 cm long with inner and outer diameters of 19 and 32 cm respectively. Notably, no electrical connections were made between the corresponding 8Tx/Rx and the 32Rx arrays, allowing for the receive arrays to be removable for service. The 8Tx/Rx_i_ array (Figure 1A,C) had two modifications vs. the 8Tx/Rx_e_ array (Figure 1D,F), the coil diameter at the inferior end of the anterior loop was increased by 5 cm, allowing for the presence of animal support tubes, and the housing was shortened to 32 cm, removing all unnecessary length while the resonant structure remained the same.

All elements in both 8Tx/Rx arrays were tuned to 447 MHz (^1^H Larmor frequency at 10.5T) using twelve lumped element capacitor (1111 E-series, Johanson Technology, Camarillo, CA, USA) locations distributed around each loop (Supplemental Figure S1A). Balanced capacitive networks (100C Series, American Technical Ceramics, Huntington Station, NY, USA) were used to impedance match each coil to the 50 Ω MR system interface. Electrically-shortened “bazooka” baluns^29^ were implemented along each coaxial cable to reduce common mode currents.

All bench measurements were recorded with a 16-channel vector network analyzer (ZNBT series, Rhode & Schwarz, Munich, Germany) while loaded with anatomically-shaped phantoms. The in vivo head-shaped and ex vivo brain-shaped phantoms (Supplemental Figure S2) were generated from previously acquired 3D Gradient Echo (GRE) data of a 7 year old 85^th^ percentile male rhesus macaque (Macaca mulatta). Both phantoms were filled with a tissue-mimicking polyvinylpyrrolidone (PVP) solution (PVP, 651.1 g/L; NaCl, 17.83 g/L; NiCl_2_-6H_2_O, 0.48 g/L; in deionized water).^30^ The conductivity and relative permittivity of the PVP solution were targeted to be 0.66 S/m and 50, respectively, at 447 MHz.^31^ Using a dielectric probe (DAKS 12, Schmid & Partner Engineering AG, Zurich, Switzerland), both the conductivity and relative permittivity were measured to be 0.64 S/m and 51.5, respectively, at 447 MHz.

Both 8Tx/Rx arrays utilized an external MR system interface^25^ which included an 8-way Wilkinson power divider^32^ with properly phased cables to allow for circular polarized (CP) excitation from the single transmit port, and eight transmit/receive (T/R) switches utilizing low noise amplifiers (SPF5122Z; Qorvo, Greensboro, NC, USA).

#### 2.1.2 32-channel In vivo Receive Array

The 32-channel in vivo receive only array (32Rx_i_) was designed to fit onto an NHP head-conformal former. The inner surface of this former measured ∼13 cm (anterior–posterior) by ∼11 cm (left–right) by ∼12 cm (superior-inferior) with a wall thickness of 2.5 mm. The 32Rx_i_ elements were constructed directly on the outer surface of the former and critically overlapped to reduce coupling.^33^ The 32Rx_i_ array (Figure 1B) was a combination of 31 loop coils and one dipole element (Figure 2A and B) positioned at the anterior-superior location, bisecting the opening above the eyes and snout, and extending up into the transverse plane. The 31 loop elements fully covered the typical radial areas (perpendicular to Z-axis) of the NHP head, while the dipole element was introduced to help increase sensitivity in the anterior-superior region (frontal lobe).^27^

**Figure 2:**
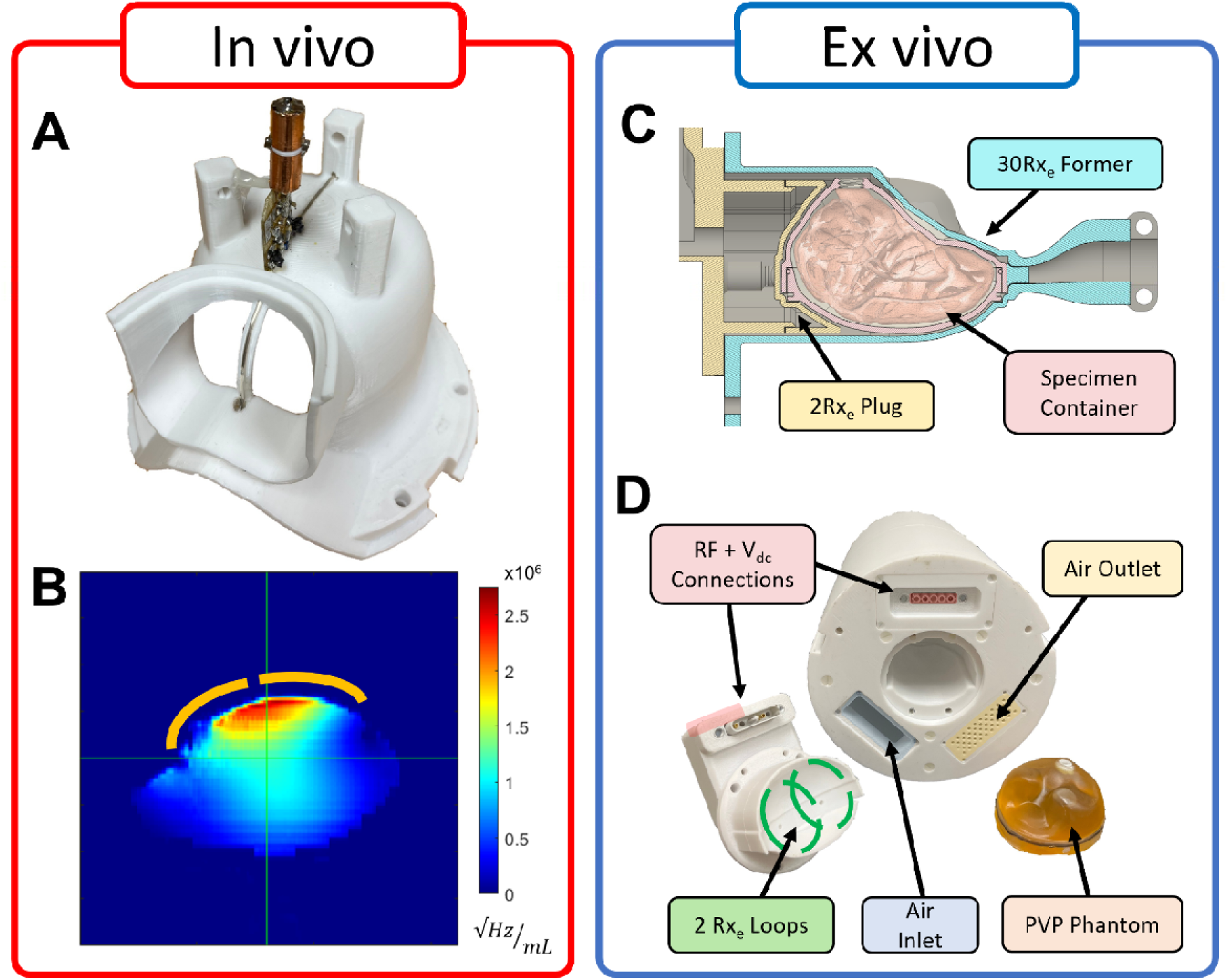
(A) Position of dipole Rx element of the 32Rx_i_ array, and (B) the resulting SNR profile from the dipole element, the arcs shown in (B) are approximate location of the dipole Rx element relative to the phantom. (C) Cross-section view of ex vivo Rx former showing the specimen fully captivated by Rx coils. (D) 2Rx plug details of the ex vivo array showing the electrical connections *(Red)* and coil placement *(Green)* within the plug, as well as, forced-air

The 31 loop elements were fabricated using 18 AWG silver-plated copper wire (MWS Wire Industries, Westlake Village, CA, USA) and ranged from 3.5 to 5 cm in diameter, with the majority being 4 cm, and formed to fit the organic shape of the former. Tuning and matching of each loop element was achieved with ceramic capacitors (1111 series, Knowles Corp, Itasca, IL, USA) located at the feed point along with a similar ceramic capacitor located along the coil conductor opposite the feed point (Supplemental Figure S1B). Passive (UM9989, Microsemi, Lowell, MA, USA) and active PIN diode (MA4P1250NM-1072T, MACOM, Lowell, MA, USA) circuitry was implemented for both coil detuning and preamp protection. All 32Rx elements were noise matched to low input impedance preamplifiers (WMA447D, WanTcom, Chanhassen, MN, USA) via semi-rigid coax cables (UT-047C-TP-LL, Micro-Coax Inc., Pottstown, PA, USA) having one or more cable traps to reduce interactions within the 32Rx_i_ array as well as with the 8Tx/Rx_i_ array. A second preamplifier protection circuit was located at the input port of the preamplifier to ensure robustness of the 32Rx_i_ array during demanding acquisitions.

The dipole antenna Rx element (Figure 2A) was fabricated from 12 AWG. silver-plated copper wire (MWS Wire Industries, Westlake Village, CA, USA). Both quarter-wavelength dipole legs were reactively shortened using air-core inductors (Midi Spring series, CoilCraft Inc., Cary, IL, USA) placed near the matching location (Supplemental Figure S1C). Using similar ceramic capacitors and diodes as the loop elements, these antenna-shortening inductors were also utilized for active and passive detuning, minimizing any interactions with the 8TxRx_i_ array. All low frequency control signals were filtered with ceramic chip inductors (1008CS series, CoilCraft Inc, Cary, IL, USA) and ceramic chip capacitors (0603 Flexicap, & 1111 series, Knowles Corp, Itasca, IL, USA) to minimize any unwanted noise within the coil arrays.

#### 2.1.3 32-channel Ex vivo Receive Array

Using a similar design and components to the 32Rx_i_ array, the 32-channel ex vivo receive only array (32Rx_e_) was designed to fit on an ex vivo NHP brain-conformal former (Figure 1E). The inner surface of this former—designed to accommodate the specimen holder detailed later herein—measured ∼5.5 cm (anterior–posterior) by ∼7.0 cm (left–right) by ∼8.5 (superior-inferior) with a wall thickness of 2.5 mm; it should be noted that the anatomical orientations listed correspond to head-first-supine positioning, though here the ex vivo brain is rotated 90° about the x-axis, resulting the in the brainstem pointing up, and frontal cortex aligned with the main magnetic field B_0_. The housing was open on the inferior end (Figures 1F) to allow for specimen placement. Figures 1E shows thirty of the 32Rx_e_ array elements constructed directly on the outer surface of the former using the same loop design and materials as the 32Rx_i_ array previously described. Notably, no dipole elements were used in the construction of the 32Rx_e_ array. Instead, a removable plug (Figures 1F) containing two of the 32Rx_e_ array elements was fabricated to phsyically captivate and fully circumscribe the imaging volume with coil elements. The 32Rx_e_ loop elements ranged from 2.5 to 4 cm in diameter, with the majority being 3.3 cm diameter and formed to fit the organic shape of the former. No passive detuning was included as the coil assembly was designed for use with ex vivo samples only. Similar to the 32Rx_i_ array, all 32Rx_e_ elements were noise matched to the same low input impedance preamplifiers, utilized the same semi-rigid coax cables and cable traps to reduce interactions within the 8TxRx_i_ array, and included the second preamplifier protection circuit located at the input port of the preamplifier.

### 2.2 Ex vivo Specimen Containers

Figure 2C and D shows custom designed containers developed to accurately and repeatedly position either the phantom load or ex vivo brain samples within the 32Rx_e_ coil assembly. The two-piece, brain-conformal containers (Supplemental Figure S2 – bottom panel) were 3D printed from a liquid resin using a stereolithography (SLA) process (Formlabs, Somerville, MA, USA). The container size and shape were designed to accommodate both hemispheres and cerebellum of various macaque species (mullata, fascicularis, et al.); the complex contours were derived from a previously acquired MRI dataset described above. The resulting container had an internal volume of 110 mL, a 225 mm circumference, and a 2.5mm thick wall thickness. All brain specimens were placed within the 2-piece containers and fixated with formalin solution (Sigma-Aldrich, St. Louis, MO, USA) prior to experiment. A 1.5 mm profile Buna-N rubber O-ring (McMaster-Carr, Chicago, IL, USA) positioned at the joint between top and bottom halves, as well as, PTFE thread sealant at the fill port were used to ensure a good seal.

### 2.3 Data Acquisition

All 10.5T data were acquired on a 10.5 Tesla Siemens MAGNETOM 10.5T Plus; (Siemens Healthineers, Erlangen, Germany) console interfaced to an 88 cm bore 10.5T magnet (Agilent Technologies, Oxford, UK) fitted with a Siemens SC72D gradient coil providing 70 mT/m maximum amplitude and 200 T/m/s slew rate. RF excitation was provided by combining eight 2 kW RF power amplifiers (Stolberg HF-Technik AG, Stolberg, Germany) into a single source, while signal reception utilized 40 of the 128 available receive channels developed in house using Siemens components.

#### 2.3.1 Data for B_1_ ^+^

Individual complex B_1_^+^ maps for each 8Tx/40Rx array were acquired using the fast relative B_1_^+^ mapping technique^34^ scaled utilizing the absolute B_1_^+^ maps acquired by actual flip-angle imaging (AFI).^35^ B1 mapping parameters of TR_1_ = 21 ms, TR_2_ = 121 ms, TE = 3 ms, flip angle = 60°, voxel size = 2.0×2.0×5.0 mm^3^.

#### 2.3.2 Data for SNR

Images for SNR calculations were obtained with a GRE sequence with NHP parameters of TR = 10000 ms, TE = 3.48 ms, flip angle = 90°, voxel size = 1.0×1.0×2.0 mm^3^, 80 interleaved axial slices covering the entire volume. Identical noise images were acquired without an RF excitation pulse and TR = 568 ms. SNR was calculated following Ugurbil, et al.^36^

#### 2.3.3 In vivo Imaging

Experimental procedures were carried out in accordance with the University of Minnesota Institutional Animal Care and Use Committee and the National Institute of Health standards for the care and use of NHPs. All subjects were fed ad libitum and single-housed within a light and temperature-controlled colony room. Animals had access to ad lib water.

On scanning days, anesthesia was first induced by intramuscular injection of atropine (0.5 mg/kg), ketamine hydrochloride (7.5 mg/kg), and dexmedetomidine (13 µg/kg). Initial anesthesia was maintained using 1.0%–2% isoflurane mixed with oxygen. For functional imaging, the isoflurane level was lowered to <1%. Each animal was wrapped in warm packs to maintain body temperature. A circulating water bath was used to provide additional heat. A ventilator was used to prevent atelectasis of the lungs, and to regulate CO_2_ levels. The animals were observed continuously, with vital signs and depth of anesthesia monitored and recorded at 15-min intervals. Rectal temperature (∼99.6 °F), respiration (10–15 breaths/min), end-tidal CO_2_ (25–40), electrocardiogram (70–150 bpm), and oxygen saturation (>90%) were monitored using an MRI compatible monitor (IRADIMED 3880 MRI Monitor, Orlando, FL, USA).

Anatomical images were obtained with a T2-weighted Turbo Spin Echo (TSE) sequence (TRLJ=LJ3000LJms, TELJ=LJ100LJms, flip angleLJ=LJ90°, voxel size = 0.5 mm iso, matrix sizeLJ=LJ260×320, iPATLJ=LJ2, turbo factor = 96, bandwidthLJ=LJ504LJHz/pixel), and SWI was acquired using a 3D-GRE sequence (TRLJ=LJ22LJms, TELJ=LJ14LJms, flip angleLJ=LJ10°, voxel size = 0.1×LJ0.1×0.5 mm, matrix sizeLJ=LJ440×732, slices = 80, iPATLJ=LJ3, bandwidthLJ=LJ190LJHz/pixel).

2D Gradient Echo EPI (GE-EPI) images for BOLD fMRI were acquired using the following parameters (TR = 1275 ms, TE = 18.6 ms, flip angle = 60°, voxel size = 0.75 mm iso, matrix size =128×154, slices = 58, iPAT = 3, multiband = 2, partial fourier = 6/8, and bandwidth = 1202 Hz/pixel). 2D single-shot Spin Echo EPI (SE-EPI) for in vivo diffusion imaging were acquired with the following parameters (TR = 8990 ms, TE = 66 ms, flip angle = 90°, voxel size = 0.75 mm iso, matrix size =144×144, slices = 74, iPAT = 3, partial fourier = 6/8, and bandwidth = 1198 Hz/pixel).

#### 2.3.4 Ex vivo Imaging

The 8Tx/40Rx_e_ ex vivo array and specimen samples were convection cooled by forced air throughout the MR experiments to reduce the temperature rise of the electronics during long diffusion acquisitions. Air input ports (Figure 2D) were included on both the 8Tx/Rx_e_ and 32Rx_e_ housings to allow connections from the remote room-temperature forced-air source (Hon&Guan, Denver, CO, USA) which was located outside of the scan room and fed into the room through waveguides using a flexible corrugated hose.

Anatomical axial and sagittal TSE images were acquired with the following parameters (TR = 3000 ms, TE = 250 ms, voxel size = .12 mm iso, matrix size = 492×800, slices = 416, iPAT = 3, bandwidth = 202 Hz/pixel). Coronal TSE data were acquired with the following parameters (TR = 3000 ms, TE = 217 ms, voxel size = .2 mm iso, matrix size = 480×480, slices = 352, iPAT = 3, bandwidth = 336 Hz/pixel).

Diffusion-weighted steady-state free precession (DW-SSFP)^37–40^ [refs] has previously demonstrated high SNR-efficiency for ex vivo diffusion imaging at ultra-high field [refs].^41,42^ Ex vivo diffusion data were acquired using a custom DW-SSFP sequence with the following parameters (TR = 21 ms, TE = 16 ms, flip angle = 14°, voxel size = 0.4 mm iso, Diffusion Gradient Duration = 10.16 ms, Diffusion Gradient Strength = 34.7, 52 mTm^-1^, Bandwidth = 222 Hz per pixel, b_eff_ = 3200, 5600 s/mm^2^). The short TR (21 ms) was compatible with a single-line k-space readout, producing images with minimal geometric distortions.

## 3 RESULTS

### 3.1 Bench Measurements

#### 3.1.1 8-channel Transmit/Receive Arrays

Bench measurements demonstrated good tuning and matching for both 8Tx/Rx_i_ and 8Tx/Rx_e_ arrays with and without their respective 32Rx_i_ or 32Rx_e_ array insert. The return losses (coil matching values, S_ii_) for the phantom-loaded 8Rx/Tx_i_ array with the 32Rx_i_ array in place ranged from −15.1 to −29.6 dB, with an average of −20.5 dB, whereas the interelement coupling (S_ij_) ranged from −9.9 to −31.2 dB with an average of −17.0 dB; 8Tx/Rx_e_ with 32Rx_e_ demonstrated comparable values. Most of the coupling within each 8Tx/Rx arrays occured between next-nearest elements, which ranged from −10.4 to −16.1 dB, with an average of −12.3 dB, and was sufficient to permit tuning of each individual element with minimal interaction. Unloaded-to-loaded Q-ratios, evaluated as Q_u_:Q_L_, were 3:1 for the 8TxRx_i_, and 2.25:1 for 8TxRx_e,_ when measured with a decoupled probe pair.

#### 3.1.2 32-channel In vivo Receive Array

All elements in the 32Rx_i_ array were matched to a minimum of −24.5 dB or better with the head-shaped phantom, with an average S_ii_ of -35.3 dB. It was impractical to measure the complete S-parameter matrix of this 32-element array; however, sample measurements for overlapped neighboring elements demonstrated -16.3 dB coupling (S_ij_) for smaller 3.5 cm loops, and -14.2 dB for larger 5 cm loops. This degree of isolation was considered sufficient given that preamplifier decoupling further improves the isolation. Among the multiple-sized array elements, the Q_U_ to Q_L_ ratios ranged from 4.4:1 for larger coil elements to 2.4:1 for smaller coil elements. During excitation the 32Rx_i_ elements demonstrated typical active detuning of ∼32 dB, with the larger loops measuring ∼25 dB, and the smaller loops measuring ∼40 dB. The two preamplifier protection circuits provided ∼56 dB of attenuation, with ∼31 dB attributed to the circuits located at the coil match boards. The dipole element of the 32Rxi was matched to -17.7 dB and demonstrated good isolation from the 8Tx/Rx_i_ elements, with S_ij_ values ranging from −14.8 to −30.9 dB while phantom loaded, prior to the detuning circuitry being applied, which further minimized any coupling.

#### 3.1.3 32-channel Ex vivo Receive Array

All elements in the 32Rx_e_ array were matched to −24.1 dB or better with the brain-shaped phantom, with an average S_11_ of -32.2 dB. Similar to 32Rx_i_, it was impractical to measure the complete S-parameter matrix of this 32-element array. As with 32RX_i_, sample measurements for overlapped neighboring elements of 32RX_e_ demonstrated -19.5 dB coupling (S_ij_) for smaller 2.5 cm loops, and -15.2 dB for larger 4 cm loops. Among the multiple-sized array elements, the Q_U_ to Q_L_ ratios ranged from 7:1 for larger, heavily loaded coil elements, to 4:1 for smaller, lightly loaded coil elements. The 32Rx_e_ elements demonstrated typical active detuning and preamplifier protection levels similar to 32Rx_i_.

### 3.2 Experimental Results

#### 3.2.1 Noise Correlation

Figure 3 shows the experimentally measured noise correlation matrices for both the 8Tx/40Rx_i_ and 8Tx/40Rx_e_ arrays while loaded with an in vivo and ex vivo specimen. The average noise correlation for the 8Tx/40Rx_i_ array was 0.181 with a maximum observed noise correlation of 0.726. The 8Tx/40Rx_e_ array demonstrated an average noise correlation of 0.167 and a maximum observed noise correlation of 0.703.

**Figure 3:**
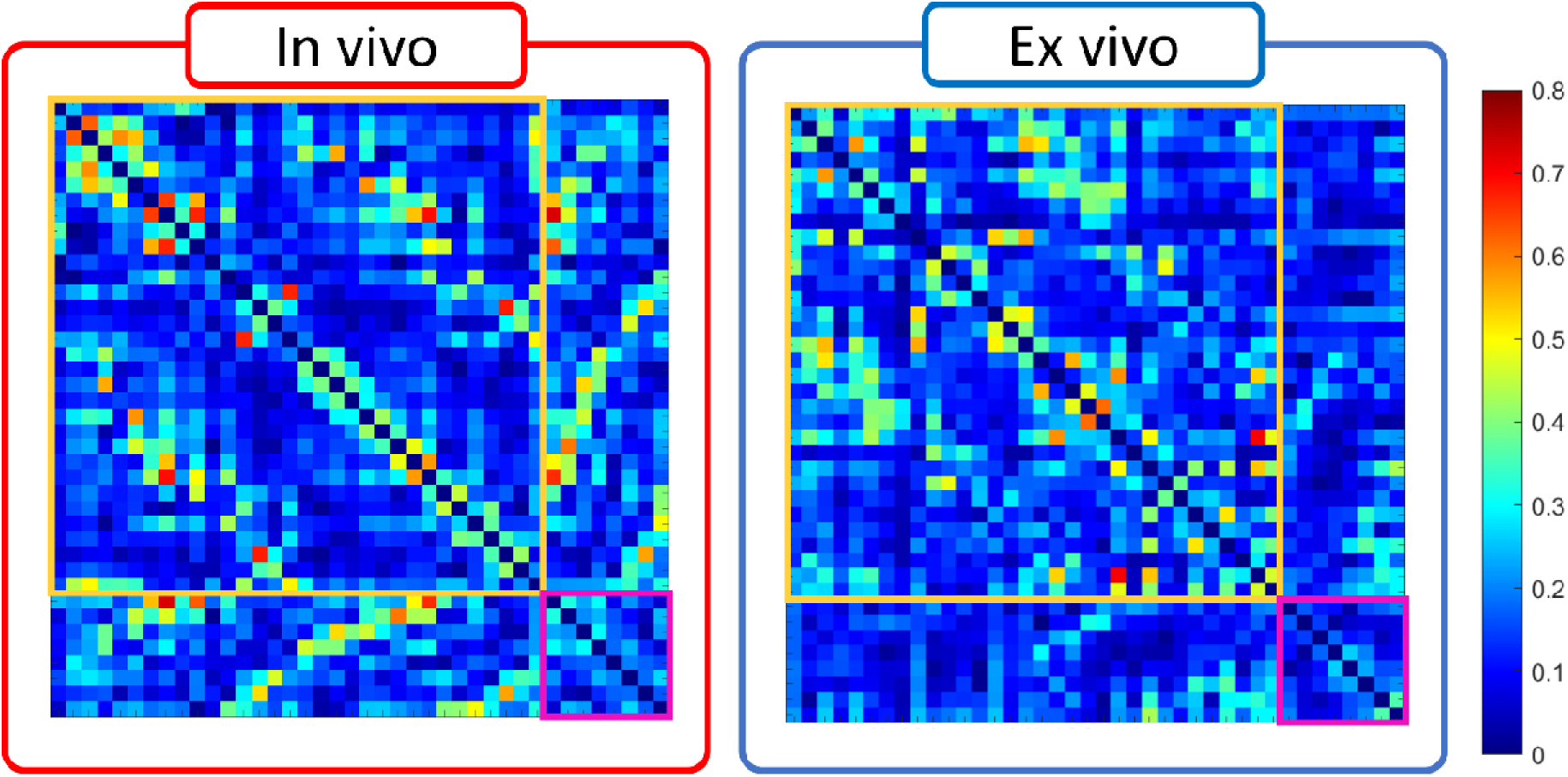
Noise correlation matrices for the 8Tx/40Rxi (A) and the 8Tx/40Rxe (B) arrays while loaded with NHP specimen. The yellow and pink boxes note the receiver (Rx) and transceiver (TR) channels, respectively.

#### 3.2.2 Phantom B ^+^

Figure 4 shows phantom-loaded B_1_^+^ maps for both coil assemblies. One can observe the higher peak B_1_^+^ within in the in vivo array due to constructive wave interactions and improved coil loading within the larger homogenous sample. The 8Tx/40Rx_i_ array produced B_1_^+^ values of 0.459 μT√w and the 8Tx/40Rx_e_ array produced 0.403 μT√w averaged across a 1 cm cubic volume located at the crosshairs indicated in Figure 4.

**Figure 4:**
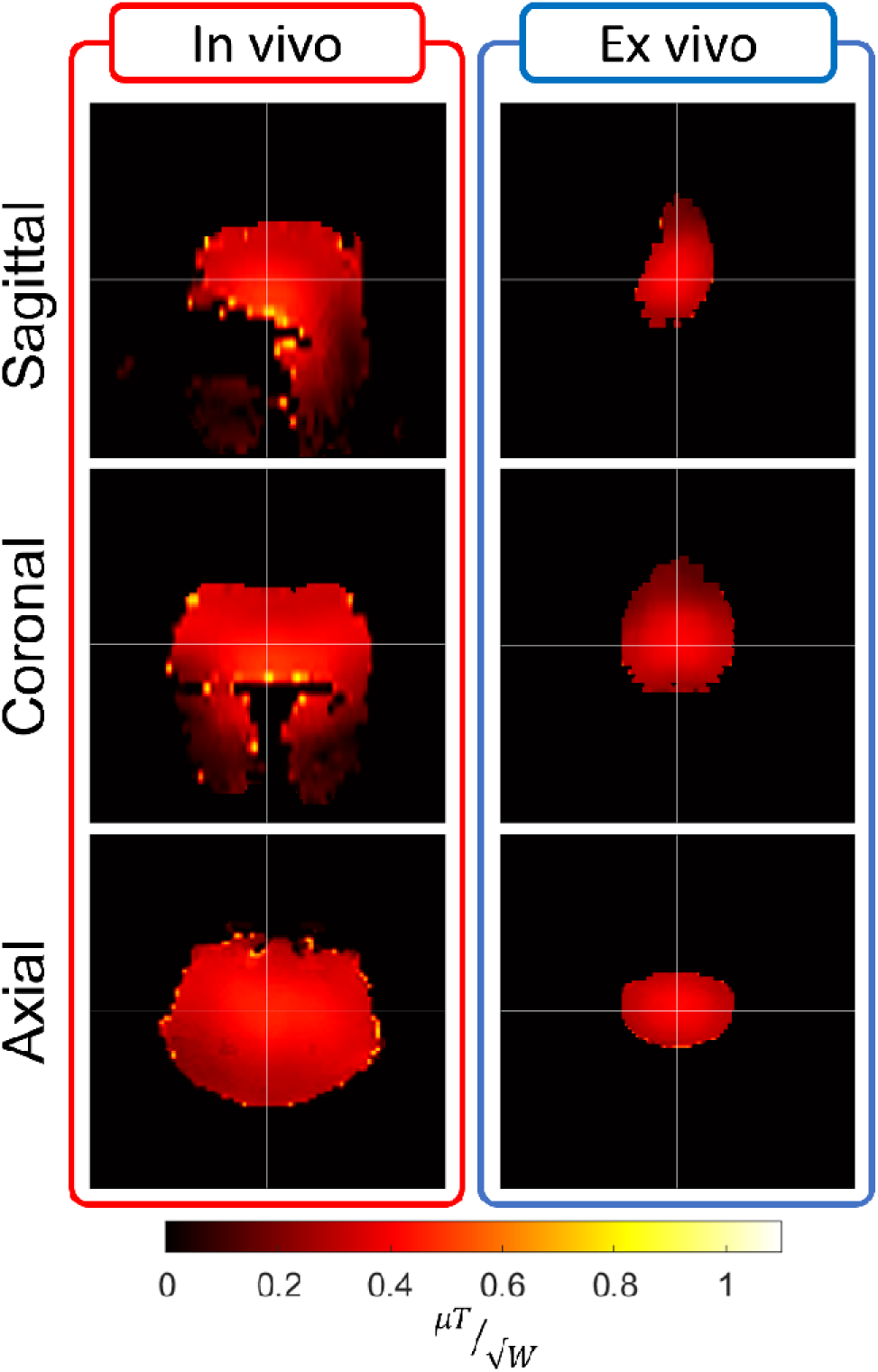
Experimentally measured B_1_^+^ maps in specimen samples, in vivo (Left), ex vivo (Right).

#### 3.2.3 SNR maps

SNR maps (Figure 5) show contributions from each subassembly (8TxRx: left column, 32Rx: center column) and their contribution to the whole (8Tx/40Rx: right column) of both 8Tx40Rx arrays in both the axial and sagittal planes. The 8Tx/40Rx_i_ array demonstrated as much as 31.1% gain in central SNR due to the transmit array being used as a transceiver, while the 8Tx40Rx_e_ showed a more modest gain of 5.5% centrally, as measured within a 1 cm cubic volume located at the crosshairs indicated in Figure 5.

**Figure 5:**
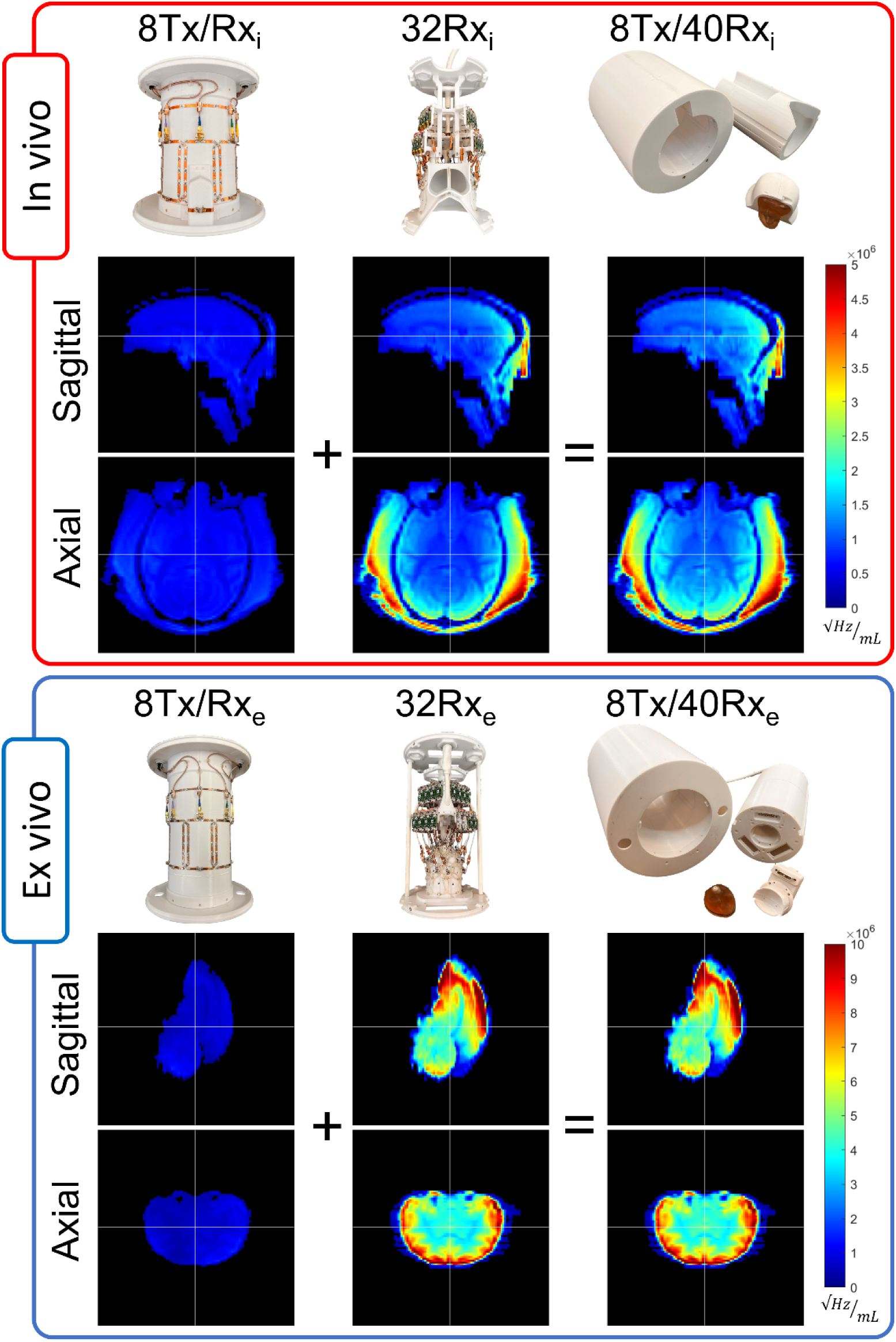
Sample SNR maps showing contributions of 8TxRx and 32Rx coil array assemblies within each 8Tx/40Rx device, in vivo (Top), ex vivo (Bottom).

The in vivo array produces high sensitivity in the temporalis muscles with relatively uniform coverage of the entire brain, while the ex vivo array shows high SNR within the entire periphery due to the conformal circumscribing 30Rx elements on the former and 2Rx within the plug *(inferior end)*.

### 3.3 In vivo & Ex vivo Imaging

#### 3.3.1 In vivo Imaging

Figure 6 (top row) shows T_2_-weighted images with 0.5 mm isotropic resolution, highlighting whole-head coverage of the 8Tx40Rx_i_ array. Figure 6 (bottom row) shows three slabs from a minimum intensity projection SWI acquisition with 0.1×0.1×0.5 mm resolution, illustrating detailed vasculature and iron rich tissue contrast throughout.

**Figure 6:**
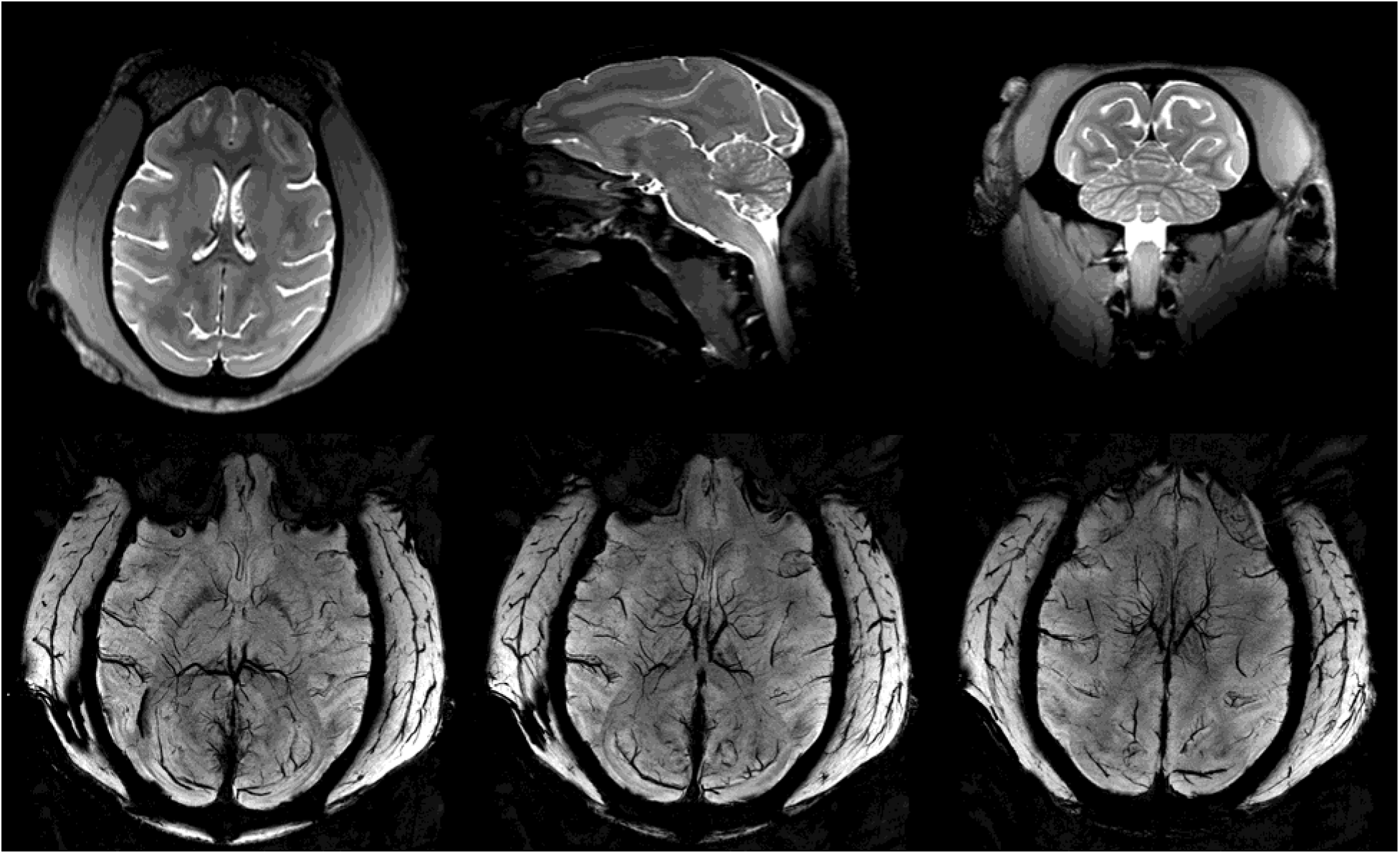
Examples of anatomical images of in vivo NHP at 10.5 Tobtained with 8-channel (Tx)/40-channel receive (Rx) array. Top row: TSE demonstrating whole head coverage. Bottom row: Minimum Intensity Projection (MIP) SWI images demonstrating deep brain iron contrast and whole-head vasculature.

Figure 7 (top row) shows 2D GE EPI with 0.75 mm isotropic resolution. Figure 7 (bottom row) shows 2D SE EPI with 0.75 mm isotropic resolution, demonstrating impressive whole-brain coverage and homogeneity.

**Figure 7:**
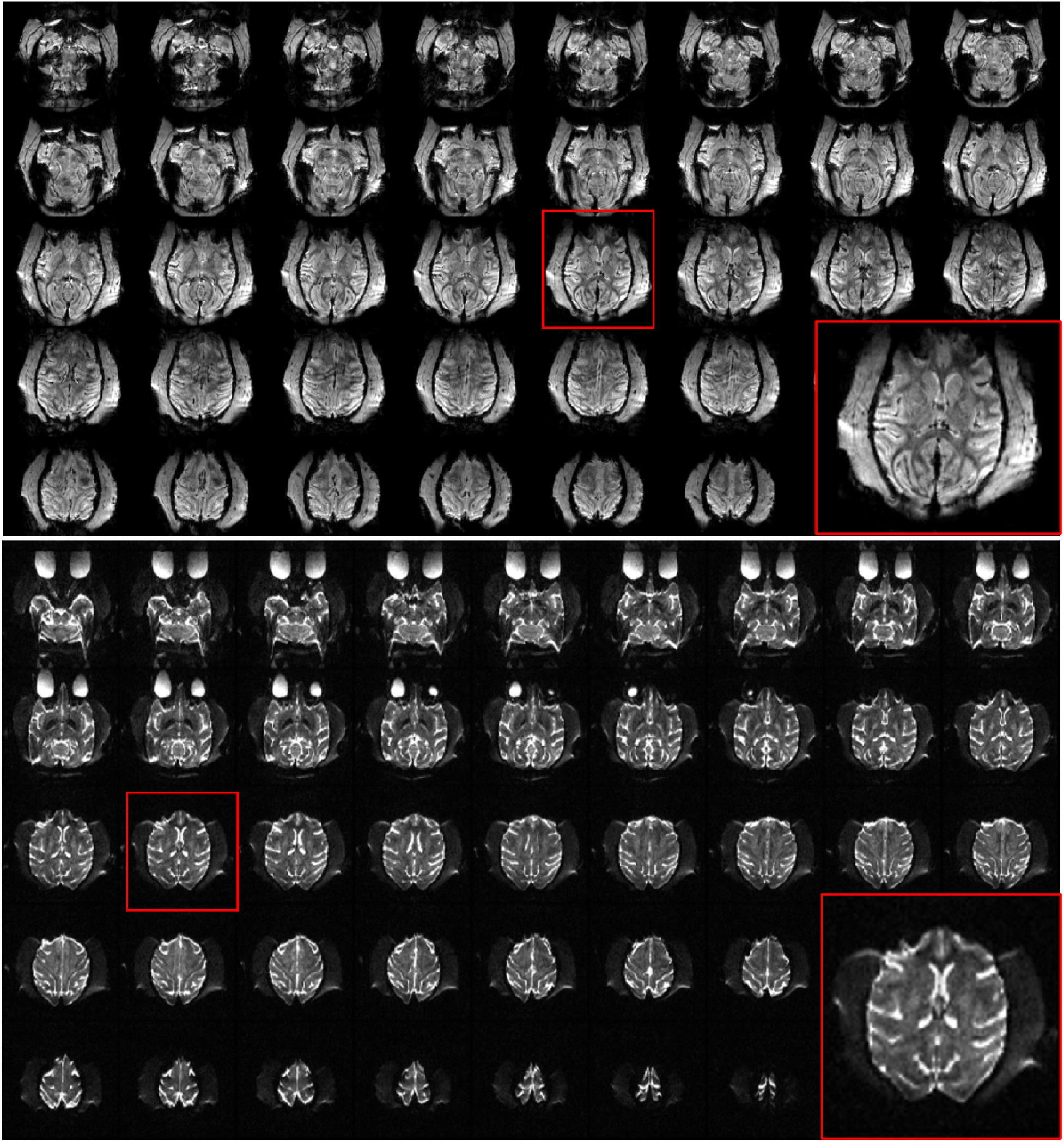
Examples of GE EPI images (top) and SE EPI *(b=0)* images (bottom), showing excellent SNR throughout the acquired whole-brain volume. A zoomed image of each series (red box) shows exceptional tissue contrast and signal homogeneity.

#### 3.3.2 Ex vivo Imaging

Figure 8a and c show uncorrected TSE images with 120 micron isotropic and 200 micron isotropic resolution respectively. Figure 8b shows DW-SSFP images acquired with two different diffusion gradient directions, yielding superb SNR and CNR with minimal geometric distortions. Orange arrows indicate well-delineated changes in the fiber orientation between the two diffusion directions.

**Figure 8:**
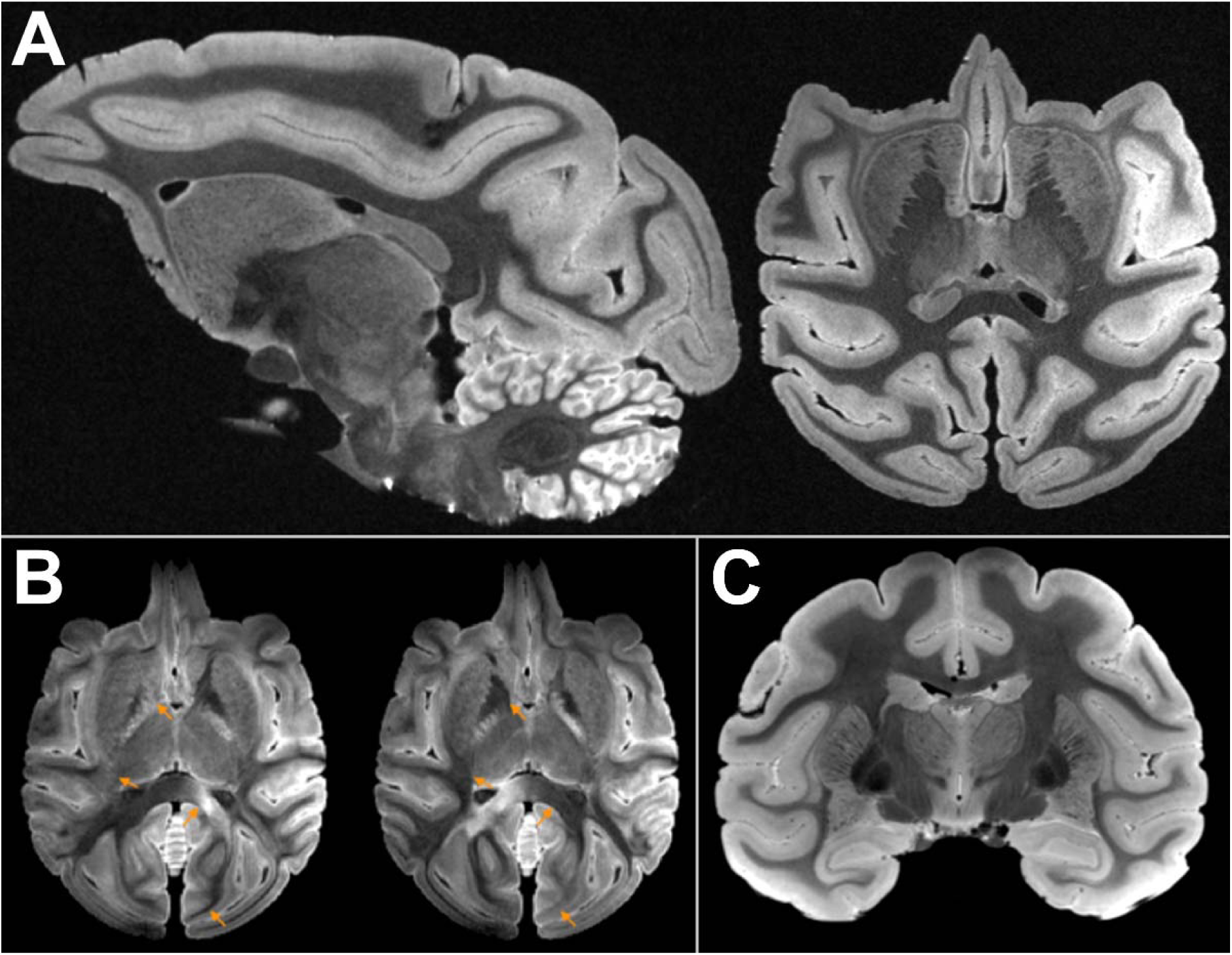
Utility examples from the 8Tx/40Rx ex vivo array: (A) Sagittal and Axial TSE at 120 micron isotropic resolution, (B) Axial DW-SSFP images at 400 micron isotropic resolution acquired with two different diffusion gradient directions (orange arrows highlight regions with notable contrast differences) at a b_eff_ = 5600 s/mm2, (C) Coronal TSE at 200 micron isotropic resolution.

Figure 9 displays the image data and resulting processed diffusion data between an in vivo and ex vivo sample. The improved spatial resolution of the ex vivo data corresponds to improved delineation of tissue anatomy and visualization of local fiber orientations, anticipated to considerably benefit future tractography investigations estimating brain connectivity.

**Figure 9:**
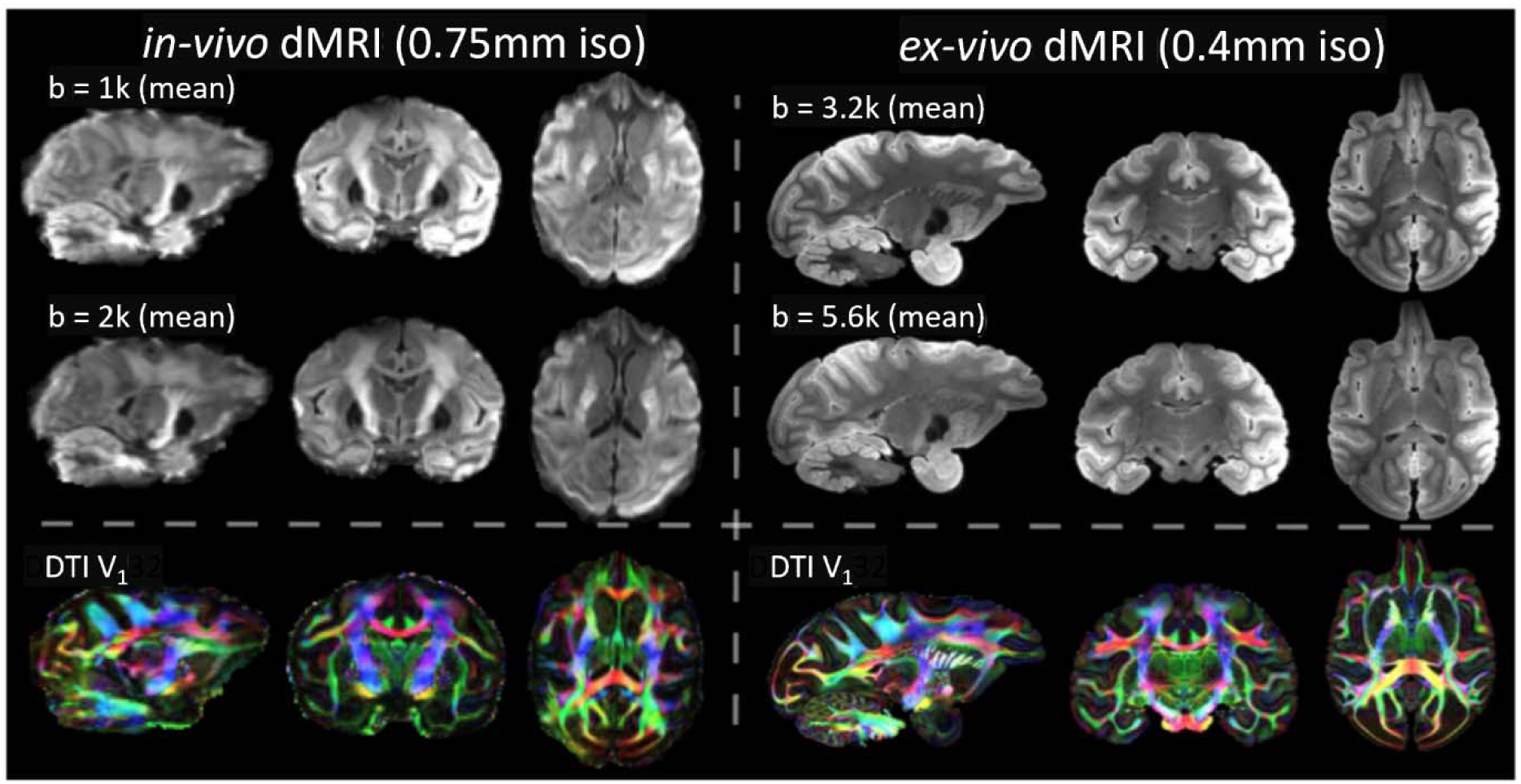
Example in vivo (Left) and ex vivo (Right) diffusion MRI datasets (averaged across all diffusion gradient directions per b-value), and resulting fractional anisotropy map color-coded by the principal fibre orientations from DTI modelling (Bottom Row). The higher spatial resolution of the ex vivo data (0.4mm isotropic) in comparison to the in vivo data (0.75mm isotropic) results in improved delineation of tissue anatomy and local fiber orientations.

## 4 Conclusions

An RF coil toolset was designed for both in vivo and ex vivo NHP brain applications at 10.5 T. The 8Tx40Rx_i_ coil array demonstrated uniform whole-head coverage for in vivo NHP. Importantly, the new coil design supported HFS positioning, which enables the harmonization of acquisition orientation and artifacts, resulting in a more direct translational approach between NHP and human imaging. The resulting orientation and large opening at the anterior of the array facilitated the use of an antenna as a receive element, and when positioned near the transverse plane, the antenna recovered some SNR in the superior and anterior brain regions where the sensitivity was reduced due to the large openings. In our previously published work^27^ we explored sensitivity profiles of various RF coil elements positioned in the transverse plane (Supplemental Figure S3), where we observed that a typical half-wavelength dipole was too long and resulted in increased coupling with adjacent Rx coils; therefore, a reactively-shortened dipole element was utilized for our application. Here we have added active detuning to the reactively-shortened dipole, which aides in decoupling the antenna receive element from the transmit array with minimal loss in deep brain sensitivity (Supplemental Figure S4). Importantly, the chosen housing dimensions can accommodate a wider range of head sizes associated with male and female monkeys of various species when compared to our previous work.^25^

The 8Tx/40Rx_e_ ex vivo coil array produced approximately 5 times higher SNR, than that of the 8Tx/40Rx_i_ array, this is expected owing to the significantly close proximity to the sample and the tightly circumscribed nature of the ex vivo array design. Higher peripheral SNR is additionally expected due to another novel feature of the design which is the lack of a circumferential” clamshell” split typical of ex vivo array architectures. Instead, the ex vivo array fully encapsulates the sample, except for an opening at the inferior end, where a plug comprising multiple receive elements can be inserted following placement of the sample within the array. This design choice not only minimizes the complexity of the design, but allows the added distance from the sample required by such “clamshell” designs to be eliminated with the associated benefits for peripheral SNR.

Parallel transmission (pTx) is a commonly used technique for producing uniform B_1_ ^+^ fields at UHF. Though the 8Tx/Rx arrays presented are capable of pTx, we found that both the in vivo and ex vivo specimen sizes were small enough to allow for CP-like excitation – which due to the associated simplicity and reduced setup time is the preferred choice. The central SNR gain (Figure 5) from utilizing the transmit array elements as receivers was beneficial for both the in vivo and ex vivo coil ensembles; however, the ex vivo improvements were more modest than we’ve previously published^3,28^, likely due to the low filling factor of the small specimen, and the high-density and proximity to the load of the tight-fitting 32Rx_e_ array.

As a result of the UHF *(10.5T)* and the various described technological innovations, both the 8Tx/40Rx_i_ and 8Tx/40Rx_e_ coil ensembles supported highest quality submillimeter resolution anatomical, functional, and diffusion imaging (Figures 6-9) within their respective in vivo and ex vivo samples. The capability of both in vivo and ex vivo MRI with the same brain specimen allows for tighter control for experimental-paradigm multistage studies examining translation between in vivo and ex vivo methodologies.

## 5 Acknowledgments

This research was funded by NIH grants UM1 NS132207, P41 EB027061, R01 EB031765-01A1, and S10 RR029672.

## Supporting Information

**SUPPLEMENTARY FIGURE S1:**
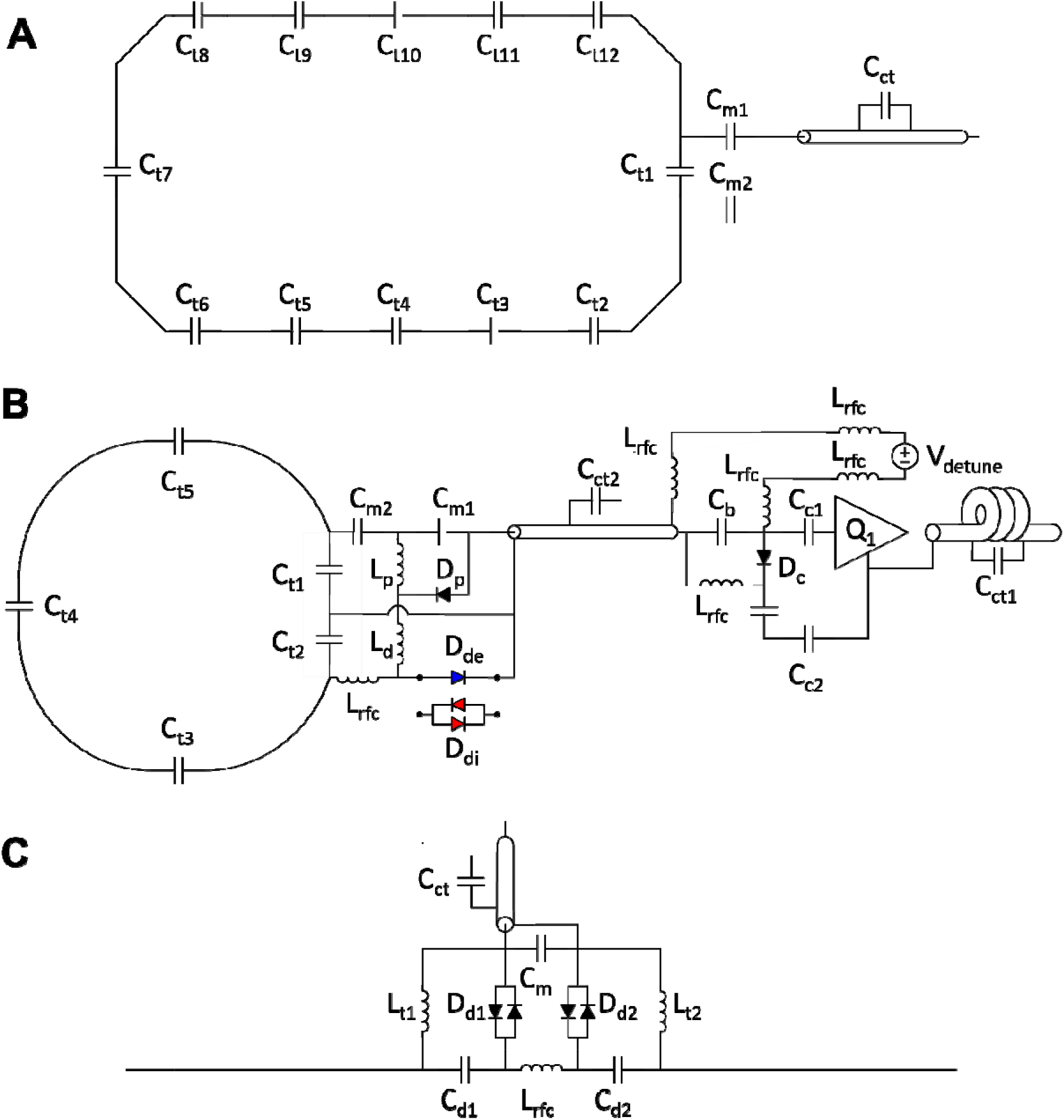
Schematic detailing (A) a single transceiver element used in both 8Tx_i_ and 8Tx_e_. (B) Single receiver element used in both in vivo (using detune diodes D_di_ - *shown in red*) and ex vivo (using diode D_de–_ *shown in blue*) arrays. And (C) actively detunable reactively-shortened dipole element used as a receiver in 8Tx/40Rx_i_.

**SUPPLEMENTARY FIGURE S2:**
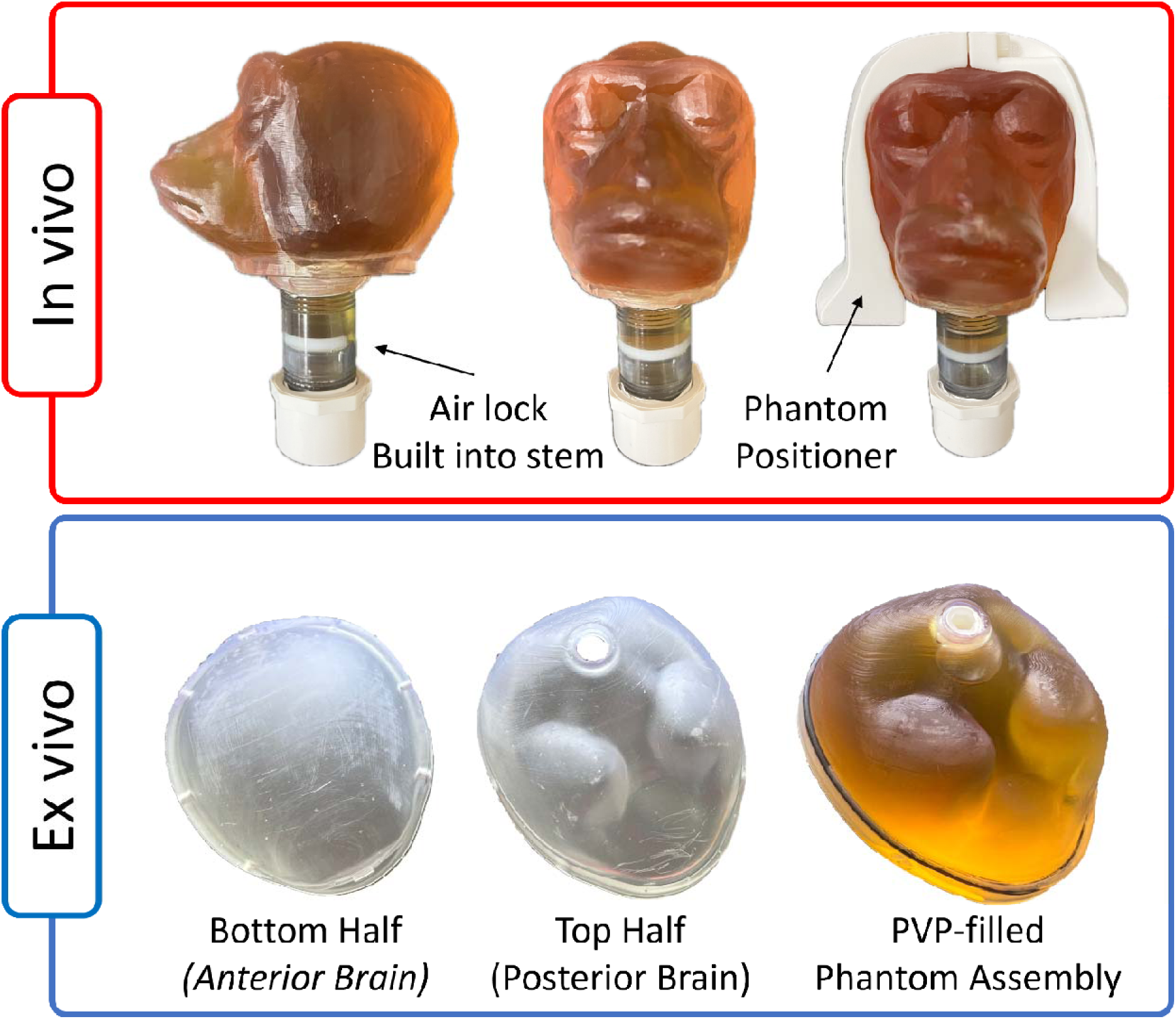
PVP-filled anatomically-shaped phantom containers for both in vivo and ex vivo applications.

**SUPPLEMENTARY FIGURE S3:**
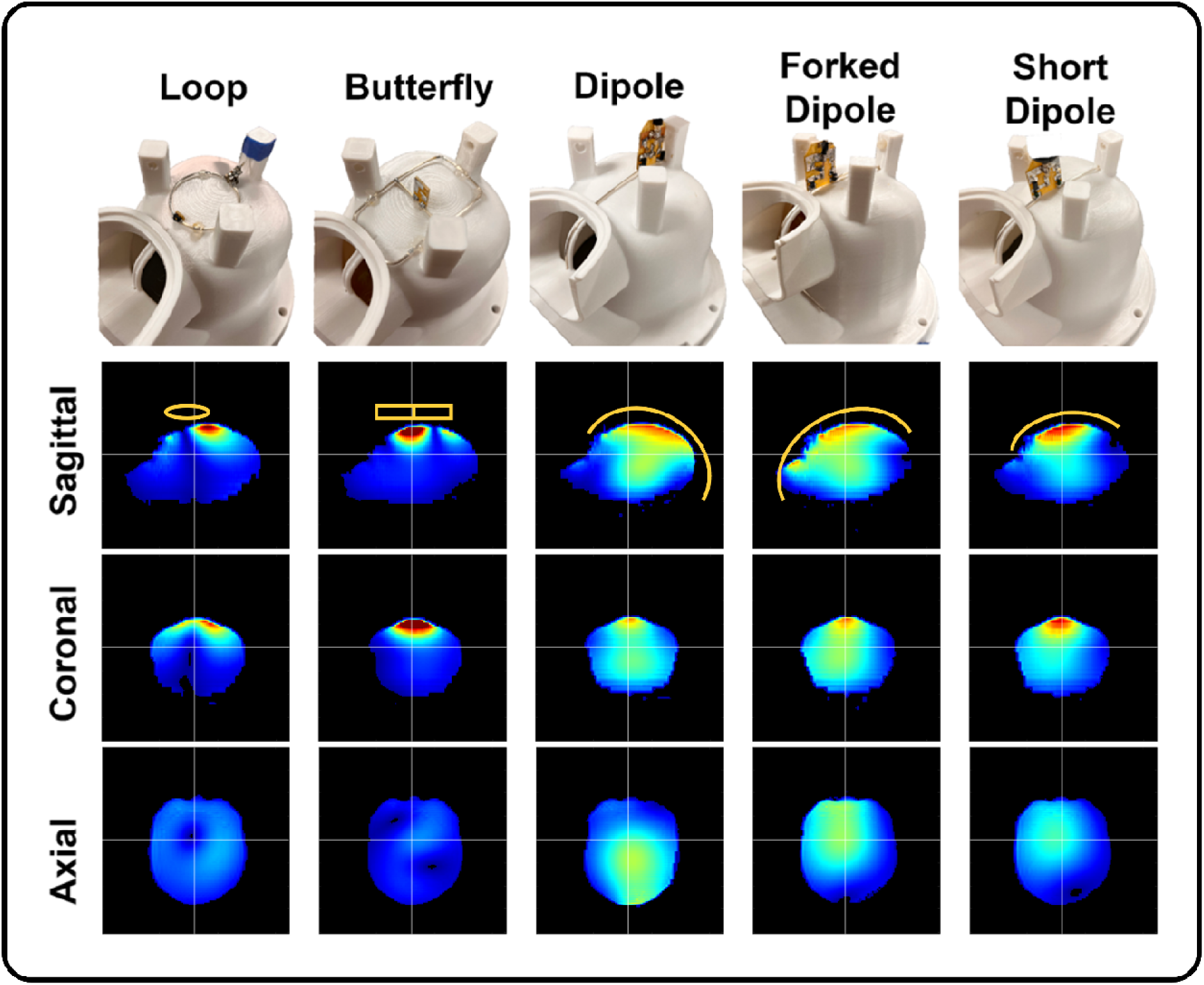
Sensitivity profiles within a PVP-filled phantom of various receive-only coil elements positioned near the X-Y plane, superior to the phantom.

**SUPPLEMENTARY FIGURE S4:**
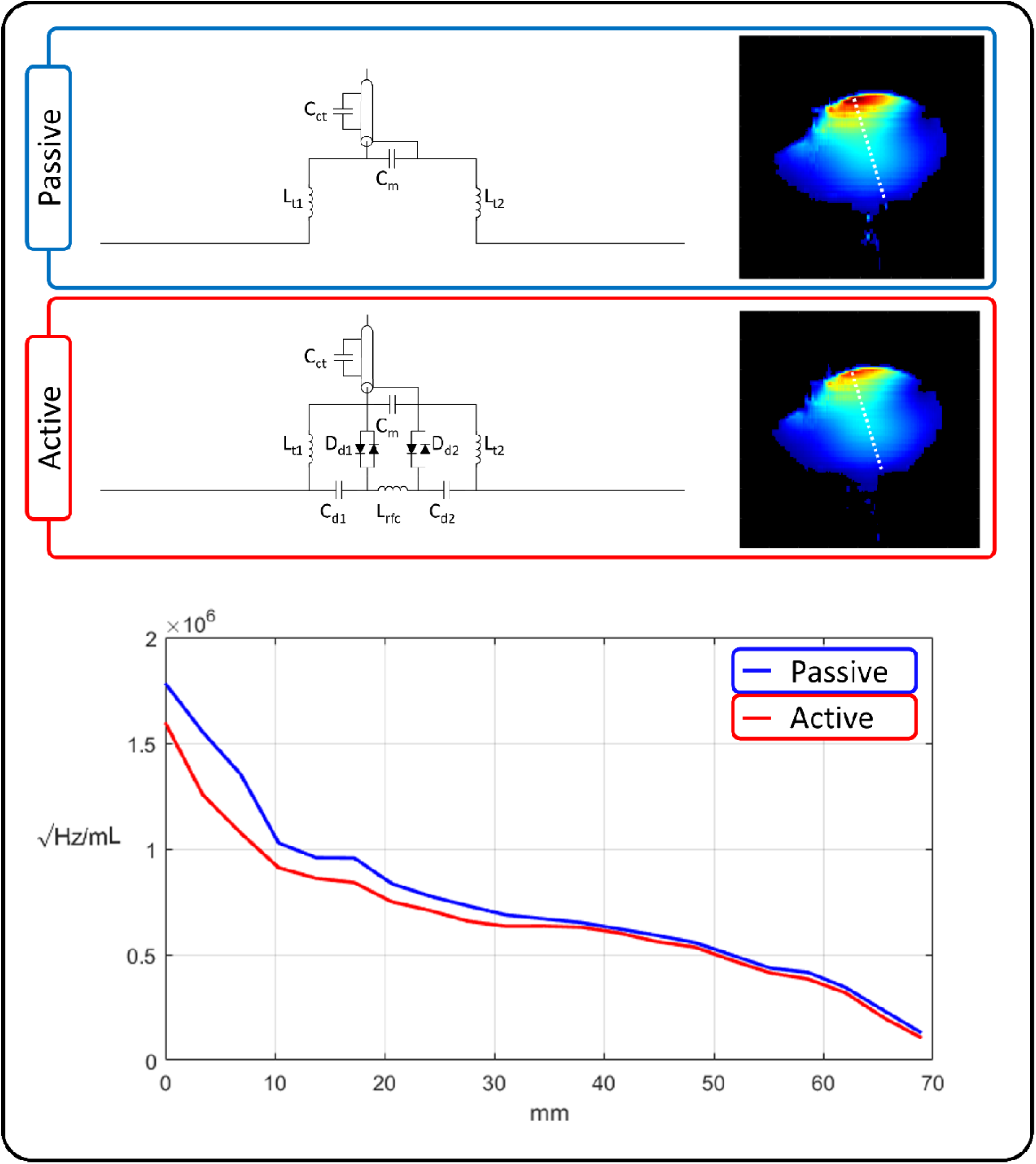
Comparison between a passive and active *(as used in 8Tx/40Rx_e_)* detunable dipole antenna. (Top) Schematics and sagittal SNR maps of passive *(blue)* and active *(red)* dipole antenna. The white dashed line indicates the path used to generate the SNR vs Depth plots (Bottom).

